# *circPCMTD1*: A protein-coding circular RNA that regulates DNA damage response in *BCR/ABL*-positive leukemias

**DOI:** 10.1101/2024.06.27.601046

**Authors:** Dimitrios Papaioannou, Amog P. Urs, Rémi Buisson, Andreas Petri, Rohan Kulkarni, Deedra Nicolet, Lauren Woodward, Chinmayee Goda, Krzysztof Mrózek, Gregory K. Behbehani, Sakari Kauppinen, Ann-Kathrin Eisfeld, Iannis Aifantis, Guramrit Singh, Adrienne M. Dorrance, Ramiro Garzon

## Abstract

Circular RNAs are a novel class of RNA transcripts, which regulate important cellular functions in health and disease. Herein, we report on the functional relevance of the *circPCMTD1* transcript in acute leukemias. In screening experiments, we found that *circPCMTD1* depletion strongly inhibited the proliferative capacity of leukemic cells with *BCR-ABL* translocations. Mass cytometry experiments identified the aberrant activation of the DNA damage response as an early downstream event of *circPCMTD1* depletion. In *in vivo* experiments, *circPCMTD1* targeting prolonged the survival of mice engrafted with leukemic blasts harboring the Philadelphia chromosome. Mechanistically, we found that *circPCMTD1* was enriched in the cytoplasm and associated with the ribosomes of the leukemic cells. We detected a cryptic open reading frame within the *circPCMTD1* sequence and found that *circPCMTD1* could generate a peptide product. The *circPCMTD*1-derived peptide interacted with proteins of the BTR complex and enhanced BTR complex formation, thereby increasing tolerance to genotoxic stress.

---

Circular RNAs (circRNAs) are a class of RNA molecules, covalently joined and characterized by the perturbed arrangement of exons that is referred to as back-splicing.^1,2^ Although their potential role in the adaptive evolution of genomes was proposed, circRNAs were initially regarded as spurious biproducts of transcription, lacking functional relevance.^3,4^ In seminal manuscripts published 30 years after their initial discovery, circRNAs were shown to play an important role in regulating mRNA translation by acting as sponges, which sequester and inhibit the function of specific microRNAs.^5,6^ Novel computational pipelines and sequencing techniques also allowed the distinction of circRNAs from spurious transcriptional products and laboratory artifacts, and enabled in-depth characterization of circRNA expression and function.^7,8^ Recent studies have delineated a plethora of mechanisms via which circRNAs regulate transcription, control translation of other mRNA transcripts and modulate cellular functions such as differentiation, proliferation and apoptosis.^9,11^ Importantly, although broadly defined as non-coding transcripts, individual circRNAs have been shown to interact with the translational machinery of the cells and generate peptides.^12,13^

In cancer, aberrant circRNA expression has been associated with cancer pathogenesis.^16,17^ Whereas specific circRNAs act as oncogenes and promote tumor growth, others act as tumor suppressors that inhibit metastasis and aggressive malignant phenotypes.^18^ Given the stability of circRNA transcripts and their resistance to exonuclease degradation, individual circRNAs have been proposed as robust biomarkers of disease persistence or progression that could inform treatment decision making.^19^ In acute leukemias, recurrent chromosomal translocations generate circRNA transcripts, which stem from the fused chromosomal junctions and which contribute to leukemogenesis.^20^ Aberrant circRNA expression has also been shown to associate with the clinical outcome of patients with cytogenetically normal AML (CN-AML).^21^ In addition, distinct circRNA signatures have been associated with the presence of recurrent prognostic mutations, such as mutated *NPM1*.^22^

In this work, we report on the functional relevance of a previously uncharacterized circRNA named *circPCMTD1* in chronic myeloid leukemia (CML) in blast phase (BP). We show that *circPCMTD1* depletion activates the DNA damage response pathway and generates a potent cell cycle block in cell lines and patient leukemia blasts. Mechanistically, *circPCMTD1* contains a complete open reading frame and interacts with ribosomes to generate a peptide. The *circPCMTD1*-derived peptide directly binds to the proteins of the BTR complex and potentiates their interaction, thereby acting as a key regulator of genotoxic stress tolerance, proliferation and cell cycle progression. In summary, we provide novel mechanistic insights into the functional relevance of circRNAs in acute leukemias and identify *circPCMTD1* as an important molecular vulnerability and a putative therapeutic target in the distinct molecular subset of myeloid malignancies with t(9;22)(q34;q11.2) translocation.

## RESULTS

### *circPCMTD1* expression associates with clinical outcome and distinct molecular features in AML

We have previously reported on the prognostic significance of circRNA expression in younger adult patients (i.e., aged <60 years) with CN-AML.^21^ The circRNA *circPCMTD1* was among the transcripts, which associated with clinical outcome in our initial study. Specifically, in our cohort of 365 younger CN-AML patients high *circPCMTD1* expression significantly associated with longer disease-free survival (*P*<0.001) and overall survival (*P*=0.001) (Supplemental Figs S1A, B, Supplemental Table S1). There was no difference in complete remission rates between patients with high and those with low *circPCMTD1* expression (*P*=0.16). With regard to clinical and molecular characteristics, patients with high *circPCMTD1* expression were more likely to have higher hemoglobin levels (*P*=0.003), lower white counts (*P*<0.001) and lower percentages of bone marrow blasts at the time of diagnosis (*P*=0.004). High *circPCMTD1* expressers were enriched among the patients who harbored *ASXL1* (*P*=0.04) and *CEBPA*-bZIPmutations (*P*=0.04). In contrast, patients with low *circPCMTD1* expression were more frequently positive for *FLT3*-ITD (*P*<0.001, Supplemental Table S1).

### *circPCMTD1* knock-down generates a strong cell cycle blockade in the molecular subset of leukemic blasts with *BCR/ABL1* translocations

To study the biologic implications of clinically relevant circRNAs, we performed functional screening experiments in a panel of myeloid leukemia cell lines. We designed LNA-modified, RNase H-recruiting, oligonucleotides (hereafter named gapmers), which targeted the back-splicing region of the candidate circRNAs transcripts and could degrade circular transcripts without affecting their linear counterparts. Following gapmer delivery, we evaluated the effect of the circRNA depletion on proliferation and cell cycle of the leukemic blasts, by performing Bromodeoxyuridine (BrdU) labelling and flow cytometry measurements.

We found that depletion of the *circPCMTD1* led to a notable decrease in the proliferative capacity and a potent G2/M blockade only in CML-BC cell lines (K-562 and LAMA-84), which harbor the t(9;22) translocation (Fig. 1A-1D, Supplemental Fig. S2A-B). In quantitative real-time PCR (qRT-PCR) experiments, we found that the effect of anti-*circPCMTD1* gapmers was specific and led to a decrease in the abundance of *circPCMTD1*, without affecting the levels of the linear *PCMTD1* (*linPCMTD1*) transcript (Fig. 1E, F). Abundance of *circPCMTD1* varied among different cell lines and LAMA-84 cells had the highest *circPCMTD1* expression (Fig. 1G). To further evaluate if the linear *PCMTD1* transcript (*linPCMTD1*) could be implicated in the strong phenotype that we observed, we designed gapmers which targeted an exon-exon junction downstream of *circPCMTD1*. The anti-*linPCMTD1* gapmers strongly depleted *linPCMTD1* without affecting *circPCMTD1* levels. *LinPCMTD1* depletion had no significant effect on the proliferation of the leukemic blasts (Supplemental Fig. S3).

**Figure 1:**
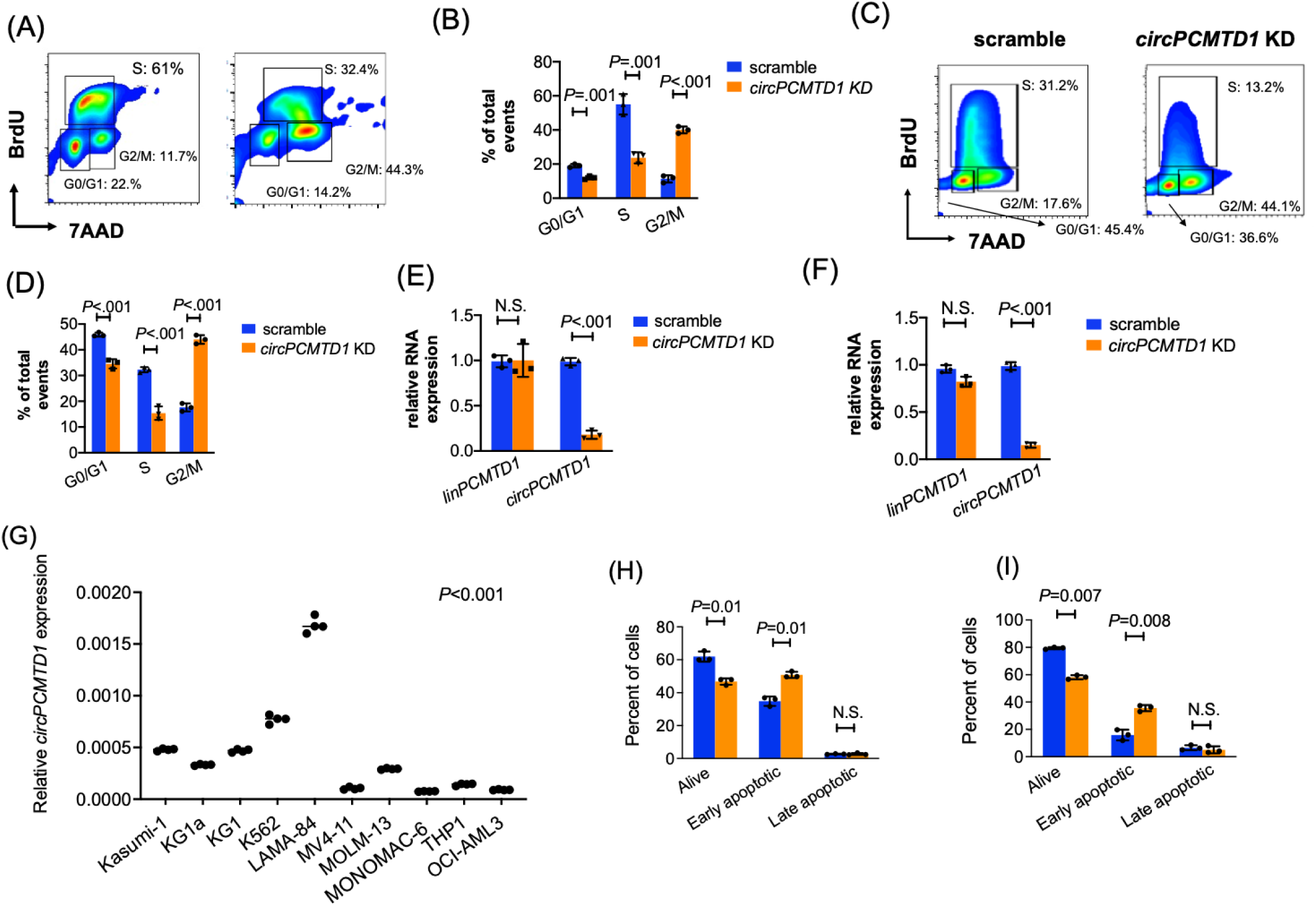
Functional impact of *circPCMTD1* depletion in BCR-ABL1-positive leukemia blasts. (A-D) BrDU labelling followed by cell cycle analysis in K-562 (A,B) and LAMA-84 (C,D) cells treated with scramble versus *circPCMTD1* KD. (B) and (D) depict the results of three independent experiments in aggregate. (E, F) *CircPCMTD1* and *linPCMTD1* expression in K-562 (E) and LAMA-84 (F) cells treated with scramble versus *circPCMTD1*-targeting gapmers. (G) Relative *circPCMTD1* expression levels in the panel of leukemia cell lines that were tested. (H, I) Percent of viable, early and late apoptotic K562 (H) and LAMA-84 (I) cells following treatment with scramble versus *circPCMTD1*-targeting gapmers. In (B), (D), (E), (F), (H) and (I), *P* values were calculated using paired two-sided t-tests. In (G) significance was tested with the Anova test. In the figures, heights of bar graphs and dot plots indicate mean values with standard deviation. Error bars indicate highest and lowest values in each population

We also evaluated the effect of *circPCMTD1* depletion in the viability of K-562 and LAMA-84 cells by Annexin/PI staining. We found that *circPCMTD1* targeting led to significant decreases in the viable fraction of the leukemic blasts (Fig. 1H, 1I).

### *circPCMTD1* knock-down increases genotoxic stress and leads to aberrant activation of the DNA damage response pathway

To gain insights and characterize the mechanistic basis of *circPCMTD1* essentiality, we first performed RNA sequencing in K-562 cells following depletion of the *circPCMTD1*. We identified approximately 150 genes which were differentially expressed following depletion of *circPCMTD1*. These included genes that are involved in cell cycle control, nuclear organization and transcriptional regulation, such as *SMARCA4*, *MACM*, *PCLAF* and *SASH1* (Fig. 2A, Supplemental Table S2). In Gene Set Enrichment analysis, we found the biological processes of rRNA processing and DNA replication-dependent chromatin function to be the notably affected by *circPCMTD1* depletion (Fig. 2B).

**Figure 2:**
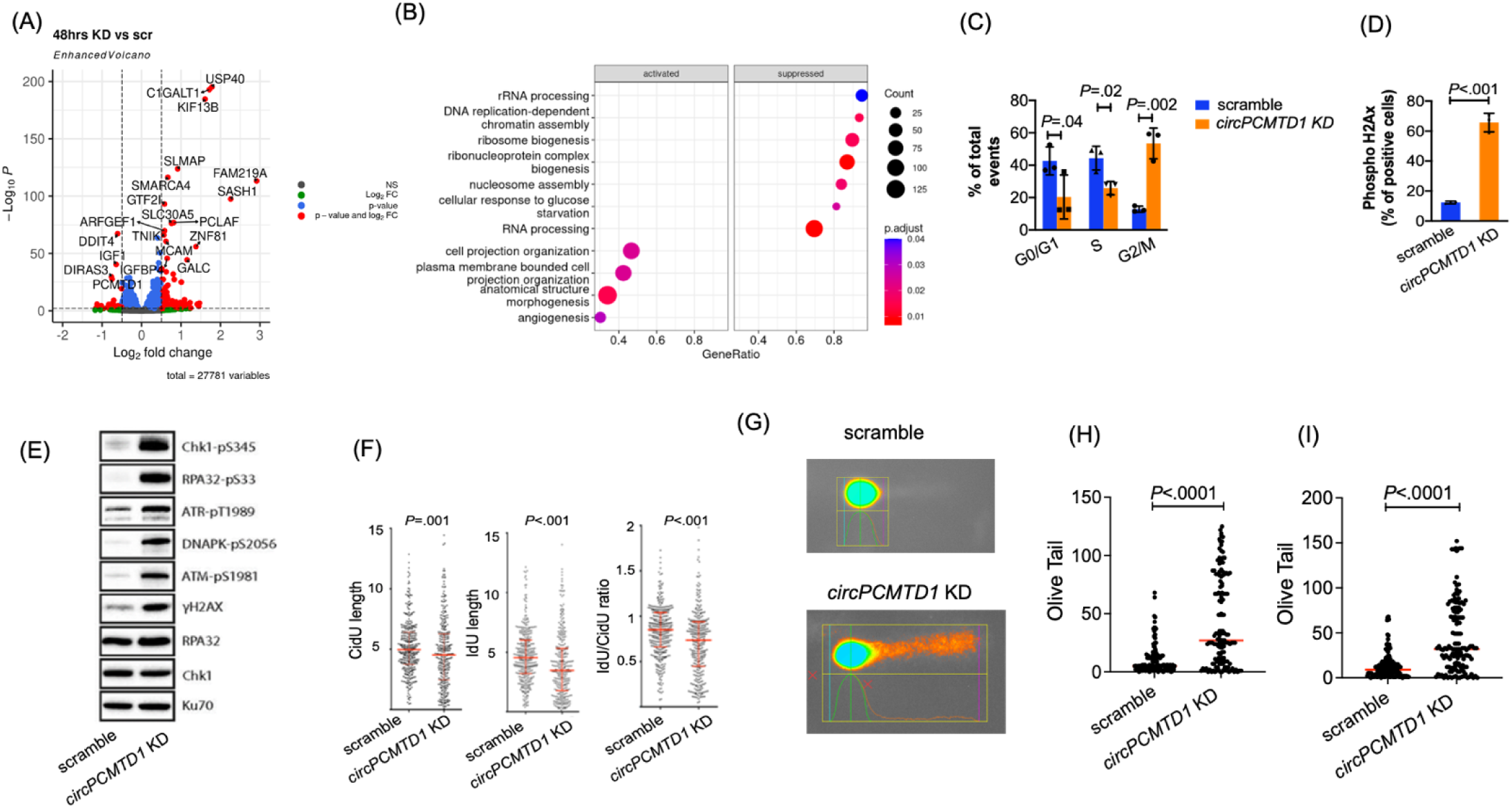
*CircPCMTD1* regulates the proliferative capacity of leukemic blasts by regulating the DNA damage response. A) Differential Gene expression and (B) Gene set enrichment analysis in K-562 cells treated with *circPCMTD1*-targeting gapmers versus controls and analyzed with total RNA sequencing. (C) Cell cycle analysis based in massive parallel flow cytometry in K-562 treated with scramble versus *circPCMTD1* KD. (D) Phospho-H2Ax levels in K562 cells treated with scramble versus *circPCMTD1* KD, as evaluated by CyTOF experiments. (E) Phospho-H2Ax (gH2Ax), CHK1, RPA32, ATR, ATM and DNA-PK in K-562 cells treated with scramble versus *circPCMTD1* KD by western blotting. Total RPA32, CHK1 and KU70 proteins are depicted as loading controls. (F) DNA fiber analysis in K-562 cells treated with scramble versus *circPCMTD1* KD. (G-I) Comet assay analyses in (G, H) K-562 and (I) LAMA-84 cells treated with scramble versus *circPCMTD1* KD. (G) depicts examples of K562 cells treated with scramble versus *circPCMTD1* KD following lysis, electrophoresis, staining and visualization per the Comet assay protocol. (H) and (I) depict results of three experiments, depicted in aggregate. In (C), (D), (F), (H) and (I) *P* values were calculated using paired two-sided t-tests. In the figures, heights of bar graphs and dot plots indicate mean values with standard deviation. Error bars indicate highest and lowest values in each population.

Furthermore, we performed massive parallel flow cytometry experiments, using the CyTOF platform. CyTOF-based cell cycle analysis of both K-562 and LAMA 84-cells validated the G2/M blockade that was generated by *circPCMTD1* depletion (Fig. 2C). In analysis of other cellular markers, we detected a significant increase in the phosphorylation of H2Ax (γH2AX) following *circPCMTD1* knock-down, which indicated aberrant activation of the DNA damage response (Fig. 2D). We confirmed the increase in γH2AX upon *circPCMTD1* depletion by western blotting (Fig. 2E) and intracellular flow cytometry (Supplemental Fig. S4). In addition, *circPCMTD1*-KD led to an increase in the phosphorylation of the CHK1, RPA32, ATR, ATM and DNA-PK proteins (Fig. 2E).

To investigate the potential role of *circPCMTD1* in the kinetics of DNA replication, we performed sequential labelling with EDU and BrdU followed by DNA fiber assays. We found that *circPCMTD1* knock-down led to a significant decrease in the length of both EdU and BrdU labelled fibers, indicating reduced capacity of the leukemic cells to initiate and maintain DNA replication upon *circPCMTD1* depletion (Fig. 2F). Furthermore, we conducted comet assays in K-562 and LAMA84 cells, following *circPCMTD1* depletion. We found that in both cell lines, depletion of *circPCMTD1* led to an increase in the fraction of DNA located in the tails of the treated cells, which suggests aberrant DNA repair response and increased rate of double-stranded DNA breaks following *circPCMTD1* depletion (Figs 2G, H). Taken together, these data underscore the aberrant DNA damage response and the significant increase in genotoxic stress that are triggered by c*ircPCMTD1* depletion.

### *circPCMTD1* expression levels are associated with disease progression in *BCR/ABL1*-positive myeloid malignancies

Our experiments revealed a strong functional effect of *circPCMTD1* in the K-562 and LAMA-84 cells and absence of a phenotype in the other leukemia cell lines that were tested. A distinctive feature of the cell lines which were sensitive to *circPCMTD1* depletion was the presence of the Philadelphia chromosome, i.e., the t(9;22) translocation. As the t(9;22) is the hallmark of chronic myelogenous leukemia, we focused on this disease and determined the relative *circPCMTD1* abundance in samples of patients at different disease stages. Specifically, we profiled samples of 17 CML patients, 9 of whom were in chronic phase, four in accelerated phase and four in blast crisis, using qRT-PCR. We found that *circPCMTD1* was more abundantly expressed in advanced disease stages (Fig. 3A).

**Figure 3:**
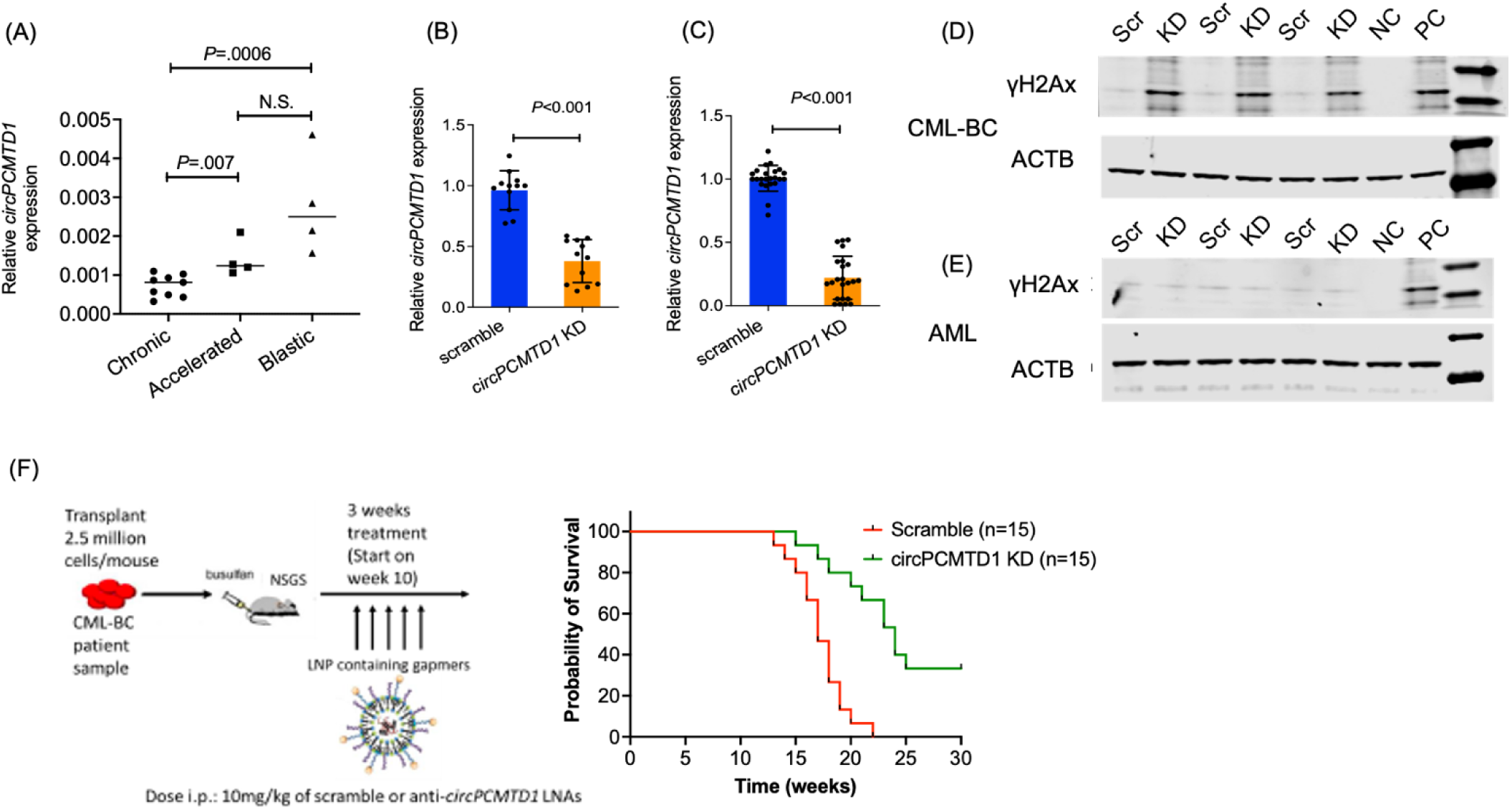
*CircPCMTD1* regulates the proliferative capacity of leukemic blasts of patients with transformed BCR-ABL+ myeloid malignancies *in vitro* and *in vivo*. (A) RNA expression levels of *circPCMTD*1 in CML patients at the chronic, accelerated and Blast phase, as evaluated by qRT-PCR. (B) *CircPCMTD1* Knockdown in leukemic blasts of CML patients in leukemic transformation. (C) *CircPCMTD1* Knockdown in leukemic blasts of AML patients. Results of 3 patients are shown in aggregate. (D, E) Phospho-H2Ax (γH2AX) staining in blasts of (D) three CML patients in blast phase and (E) three AML patients following *circPCMTD1* KD. B-actin is used as loading control. NC indicates negative control, PC indicates positive control. (F) Schematic representation of *in vivo* delivery of *circPCMTD1*-targeting oligos versus controls in mice xenografted with human CML-BP blasts and Kaplan Maier curves depicting the survival of the treated mice. In (A–C), P values were calculated using paired two-sided t-tests. Heights of dot plots and bar graphs indicate mean values with standard deviation. Error bars indicate highest and lowest values in each population. In (F), P values were calculated using the log-rank test.

### *circPCMTD1* knock-down activates DNA damage response and suppresses leukemic outgrowth of *BCR/ABL1*-positive patient blasts

To confirm our initial observations in leukemia cell lines regarding the functional relevance of *circPCMTD1*, we performed *in vitro* experiments with leukemic blasts of AML patients and CML patients in blast crisis. Although *circPCMTD1* expression was comparable and efficiently depleted in all samples that were tested (Fig. 3B, C), *circPCMTD1* knock-down had a distinct effect on BCR-ABL positive blasts. Specifically, *circPCMTD1* depletion led to an increase in the γH2Ax levels in all *BCR/ABL*-positive samples that were tested (as evaluated by western blotting). Conversely, no increase in γH2Ax levels was noted in the *de novo* AML samples (Fig. 3D, E).

To further evaluate whether *circPCMTD1* targeting could impair growth of the leukemic blasts *in vivo*, we performed experiments in immunocompromised [i.e., NOD-scid IL2Rgamma^null^ (NSG)] mice engrafted with *BCR/ABL*-positive patient blasts (collected during blast crisis). We randomly divided the xenografted animals into two cohorts, each consisting of 15 mice. On day 10 post-transplant, we began treatment of each cohort with either non-targeting controls (cohort A) or anti-*circPCMTD1* targeting gapmers (cohort B). All mice showed evidence of engraftment prior to initiation of treatment. Gapmers were packaged in cationic lipid nanoparticles, as previously described.^23^ Treatment was continued for three weeks. We detected a significant prolongation in the survival of the animals treated with *circPCMTD1*-targeting gapmers versus the cohort treated with non-targeting controls. The median survival in cohort A was approximately 20 weeks. All mice died of fulminant leukemia. In contrast, the median survival of cohort B was 26 weeks (*P=*0.002, Fig. 3F). Notably, one third of the mice in cohort B survived until the completion of the experiment and had no detectable disease at that timepoint (36 weeks of treatment). There were no notable toxicities from the treatment with cationic lipid nanoparticles and no deaths from causes other than leukemia.

### *circPCMTD1* is enriched in the cytoplasm of AML cells and associates with the ribosomes

To further characterize the mechanistic relevance of *circPCMTD1* expression in Ph+ acute leukemias, we first evaluated the subcellular localization of the *circPCMTD1* transcript. We purified fractions of chromatin, nucleoplasm and cytoplasm from K-562 cells and evaluated the abundance of *circPCMTD1* in each compartment. We used *GAPDH* (which is predominantly located in the cytoplasm and *MALAT1* (which is retained in the nucleus) as controls for the purity of each fraction. We found that *circPCMTD1* was most abundant in the cytoplasm of the leukemic blasts (Fig. 4A-C).

**Figure 4.**
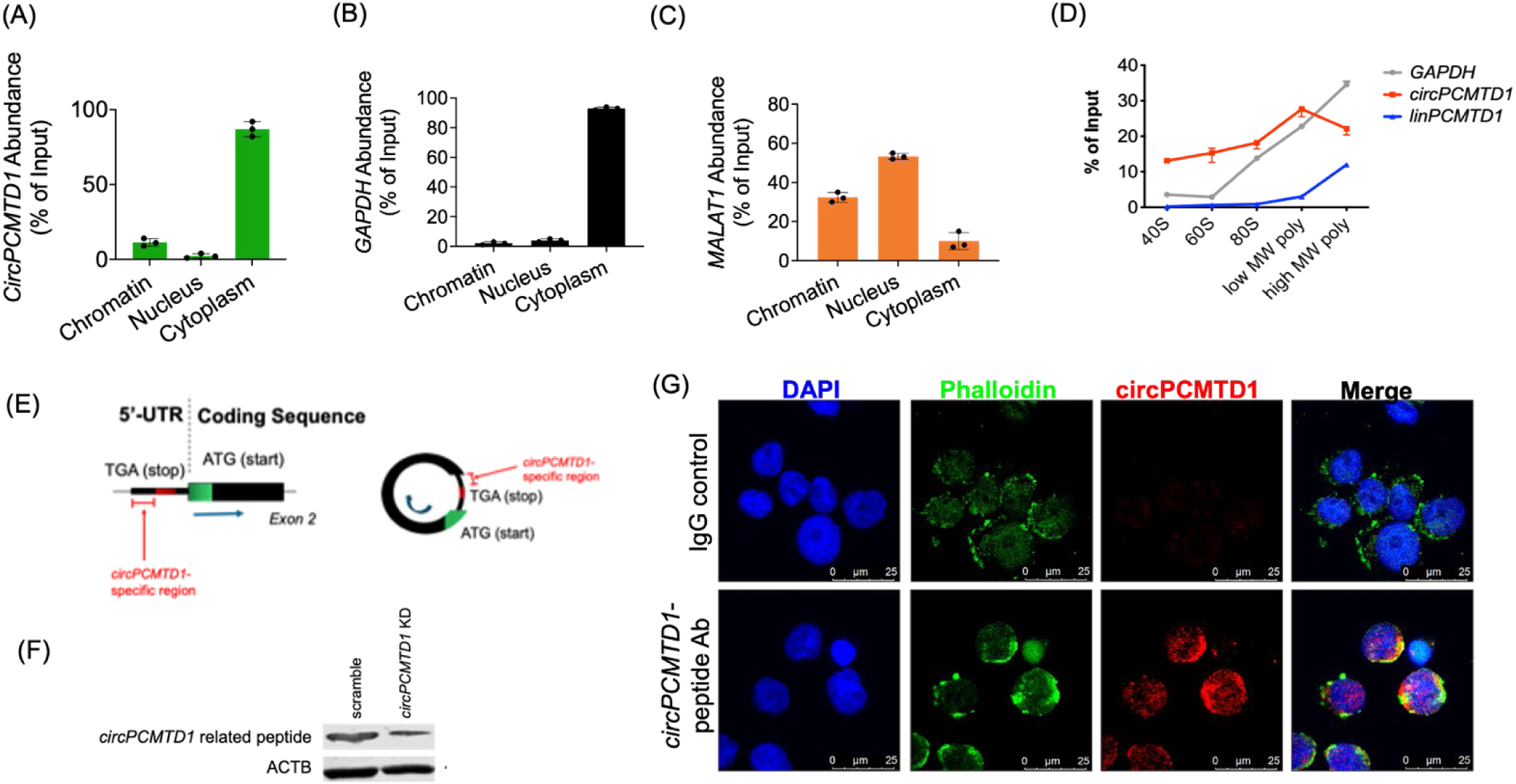
*CircPCMTD1* associates with the ribosomes of the leukemic blasts and generates a peptide. (A-C) Subcellular localization of the *circPCMTD1* transcript (A) in K-562 cells. *GAPDH* (B) and *MALAT1* (C) were used as controls (D) Targeted polysome profiling and association of the *circPCMTD1*, *GAPDH* and *linPCMTD1* transcripts with the 40S, 60S and 80S ribosomal units as well as the light- and heavy-molecular weight polysomes. (E) Schematic representation of the open reading frame contained in the *circPCMTD1* transcript. (F) Western blotting depicting the decreased levels of the *circPCMTD1*-derived peptide after *circPCMTD1*-KD in K562 cells. (G) Immunofluorescent imaging visualizing the subcellular distribution of the *circPCMTD1*-derived peptide in LAMA-84 cells. Heights of bar graphs indicate mean values with standard deviation. Error bars indicate highest and lowest values in each population.

Given its similar distribution with protein-coding genes and recent reports of circRNAs with protein-coding potential, we performed polysome profiling and evaluated the association of *circPCMTD1* with the translational machinery of the leukemic blasts. We found enrichment of the *circPCMTD1* in the light-weight fractions of the polysome complexes, which suggested a potential protein-coding capacity of *circPCMTD1*(Fig. 4D). Importantly, we found that *circPCMTD1* contained a complete open reading frame, with all functional elements of a protein-coding transcript, which was uniquely present in the circular configuration of the transcript. Specifically, we detected a stop codon in the 5-prime untranslated region of exon 2 of *PCMTD1*. In the linear configuration of the transcript this stop codon would have no functional relevance (as it is located upstream of the start codon which signals initiation of the translation). Upon circularization of exon 2, however, the stop codon became in-frame with the start codon. The cryptic open reading frame consisted of 128 codons, 24 of which would be uniquely present in a potential *circPCMTD1*-derived peptide (Fig. 4E, Supplemental Table S3).

We used the *circPCMTD1*-specific region of the predicted peptide to generate synthetic epitopes. These epitopes were used to immunize rabbits and generate an antibody against the *circPCMTD1-* derived peptide. This antibody detected a protein product of approximately 30 KD in size, which was decreased in expression after delivery of *circPCMTD1*-targeting gapmers (Fig. 4F).

We then sought to characterize the subcellular localization of the *circPCMTD1*-derived peptide. We performed immunofluorescence experiments using our custom designed antibody. In keeping with our findings which suggest a role of the *circPCMTD1*-derived peptide in DNA damage response, the majority of the detected peptide was localized in the nucleus of the leukemic blasts (Fig. 4G).

### The *circPCMTD1*-derived peptide interacts with the proteins of the BTR complex

To further elucidate the mechanistic function of the *circPCMTD1*-derived peptide, we performed immunoprecipitation experiments with our custom-designed antibody, followed by mass spectrometry analysis. In brief, we incubated nuclear lysates of K-562 cells with the antibody against the *circPCMTD1*-derived peptide or rabbit IgG control. We analyzed the eluates with tandem mass spectrometry to identify putative interacting proteins of the *circPCMTD1*-derived peptide. Based on the ratio of peptides detected in the *circPCMTD1*-derived peptide immunoprecipitate versus the control, we identified approximately 23 candidate interactors (Supplemental Table S4). Among the proteins with the highest ratio of identified peptides, we detected the BLM protein (Fig. 5A). BLM is a DNA helicase that unwinds single- and double-stranded DNA and is involved in DNA replication and repair.^25–27^ Notably germline mutations in the BLM gene generate strong clinical phenotypes (i.e., the Bloom syndrome, which is classified as a congenital bone marrow failure syndrome with a propensity to leukemia transformation).^28^ Among the remaining proteins with high ratio of identified peptides against the IgG control, we identified the TOP3A and RMI1 proteins. TOP3A is a topoisomerase, involved in preventing DNA damage during replication by facilitating DNA strand breakage by super-coiling.^29,30^ RMI1 is an essential binding partner of the TOP3A protein, that facilitates its function and maintains genome integrity.^31^ Importantly, BML, TOP3A and RMI1 interact to form the BTR complex, which is also referred to as the BLM dissolvasome, an important regulator of DNA integrity during replication.^32,33^

**Figure 5:**
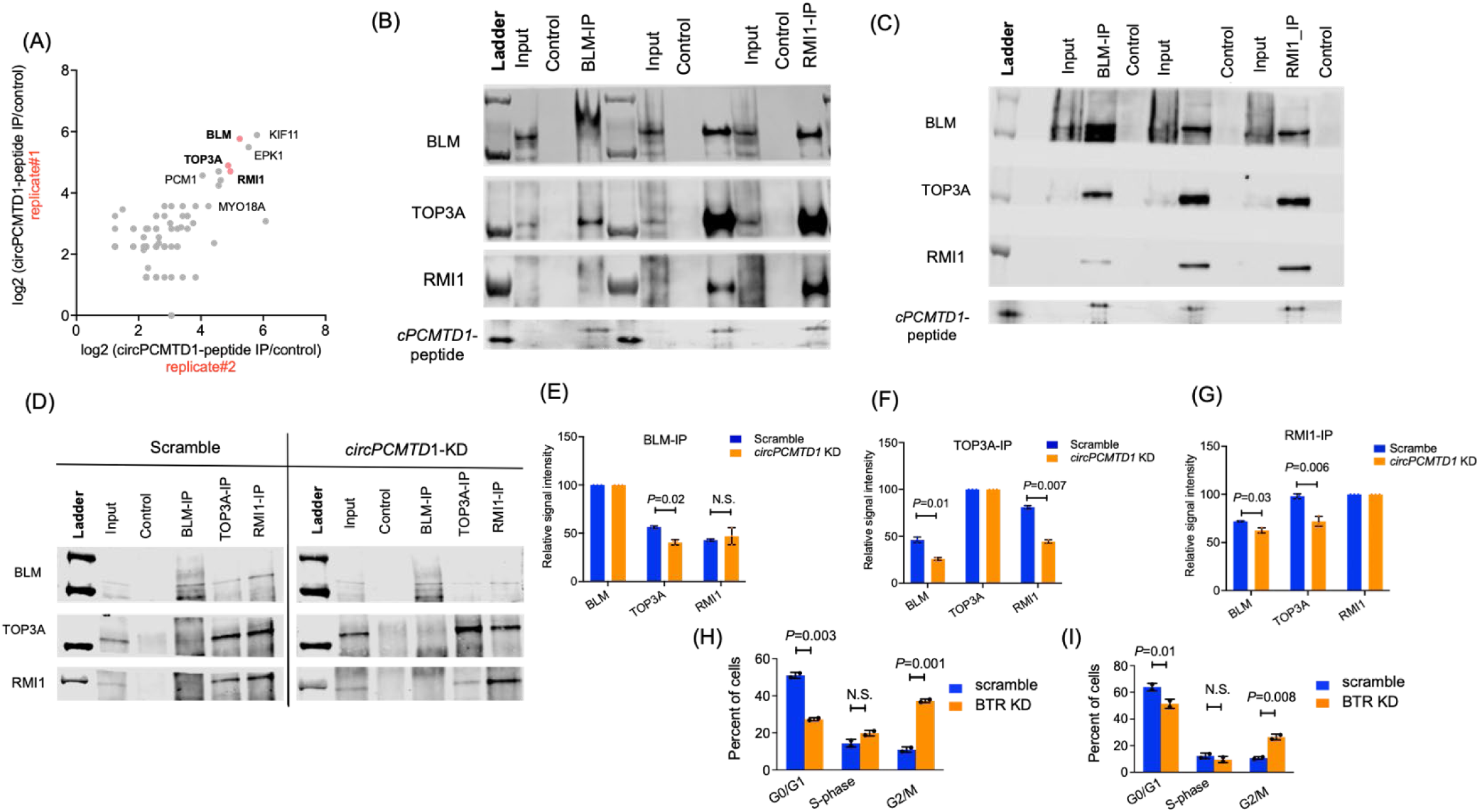
The *circPCMTD1-*derived peptide associates with the proteins of the BTR complex and affects the strength of their interaction. (A) Putative protein interactors of the *circPCMTD1*-derived peptide as identified by mass-spectrometry analysis. (B, C) Immunoprecipitations with the custom antibody against the *circPCMTD1*-derived peptide, followed by western blotting for the BLM, TOP3A, RMI1 proteins in (B) K562 and (C) LAMA-84 cells. (D) Immunoprecipitations with antibodies targeting the BLM, TOP3A, RMI1 proteins, followed by western blotting in K562 cells treated with scramble versus *circPCMTD1* KD. (E-G) Quantification of the strength of interaction among the BTR proteins following *circPCMTD1*-KD by Image J. Effect of the BTR protein complex KD on cell proliferation of (H) K562 and (I) LAMA-84 cells. In (E)-(I), P values were calculated using paired two-sided t-tests. In the figures, heights of bar graphs indicate mean values with standard deviation. Error bars indicate highest and lowest values in each population.

To confirm the interaction of the *circPCMTD1*-derived peptide with the proteins of the BTR complex, we first performed immunoprecipitations in K562 and LAMA-84 cell lines (Fig. 5 B,C). In both cell lines, immunoprecipitation of each of the BLM, TOP3A and RMI1 proteins followed by western blotting, confirmed their interaction and the interaction with the *circPCMTD1*-derived peptide (Fig. 5 B,C).

### The *circPCMTD1*-derived peptide facilitates the interaction of the BLM, TOP3A and RMI1 proteins and regulates the formation of the BTR complex

Given the interaction between the *circPCMTD1*-derived peptide and the proteins of the BTR complex, we first evaluated the effect of *circPCMTD1* depletion on the abundance of the proteins involved in the BTR formation. *circPCMTD1* knock-down had no significant effect on the levels of the BLM, TOP3A or RMI1 proteins (Supplementary Fig. S5). *CircPMTD1* depletion did however have a notable impact on the interaction of the three proteins and the formation of the BTR complex. Specifically, we found that upon *circPCMTD1* depletion the strength of interaction between the BTR complex proteins was decreased and a lesser amount of each of the interacting proteins was detected when immuno-precipitants of BLM, TOP3A and RMI1 were analyzed with western blotting (Fig. 5D-G, Supplementary Fig. S6).

To confirm that the decreased amounts of the formed BTR complex was indeed the molecular pathway driving the phenotype that was observed with *circPCMTD1* depletion, we performed knockdowns of the BLM, TOP3A and RMI1 proteins individually and combined (to concomitantly target all components of the BTR complex). While knock-downs of the individual proteins reduced the viability of leukemic blasts but had a modest effect on cell cycle progression, concomitant depletion of BLM, TOP3A and RMI1 led to the G2/M cycle blockade that we observed upon *circPCMTD1* knock-down (Fig. 5H-I, Supplementary Fig. S7).

### Inhibition of the constitutively active BCR/ABL1 kinase abrogates the association of the *circPCMTD1*-derived peptide with the BTR complex

Given the pronounced effect of *circPCMTD1* depletion in the subset of blasts with *BCR/ABL1* translocations, we wanted to evaluate the role of the constitutively active ABL1 kinase (which is the hallmark pathogenic event of *BCR/ABL1* translocation) and its down-stream signaling in the function of *circPCMTD1*. To this end we treated K562 cell with Dasatinib, an established front-line tyrosine kinase inhibitor (TKI) in sublethal doses, in order to block BCR/ABL1 signaling without causing apoptosis of the leukemic cells (i.e., 1 nM). Reduced phosphorylation of the STAT5 protein was used as a control of efficient BCR-ABL1 blockade (Fig. 6A, B). Interestingly, treatment with Dasatinib led to a decrease in the detectable levels of the *circPCMTD1*-derived peptide without affecting the expression levels of the *circPCMTD1* transcript. In addition, we evaluated the effect of TKI treatment on the interaction of the BTR complex at early timepoints, when the detected levels of the *circPCMTD1*-derived peptide were not altered. In immunoprecipitation experiments we found that TKI treatment reduced the strength of the interaction between the BLM, TOP3A and RMI1 proteins and the formation of the BTR complex (Fig. 6C-E). These results suggest a link between the aberrant BCR/ABL1 kinase activity, the abundance of the *circPCMTD1*-derived peptide and its capacity to enhance BTR formation and regulate the DNA damage response.

**Figure 6:**
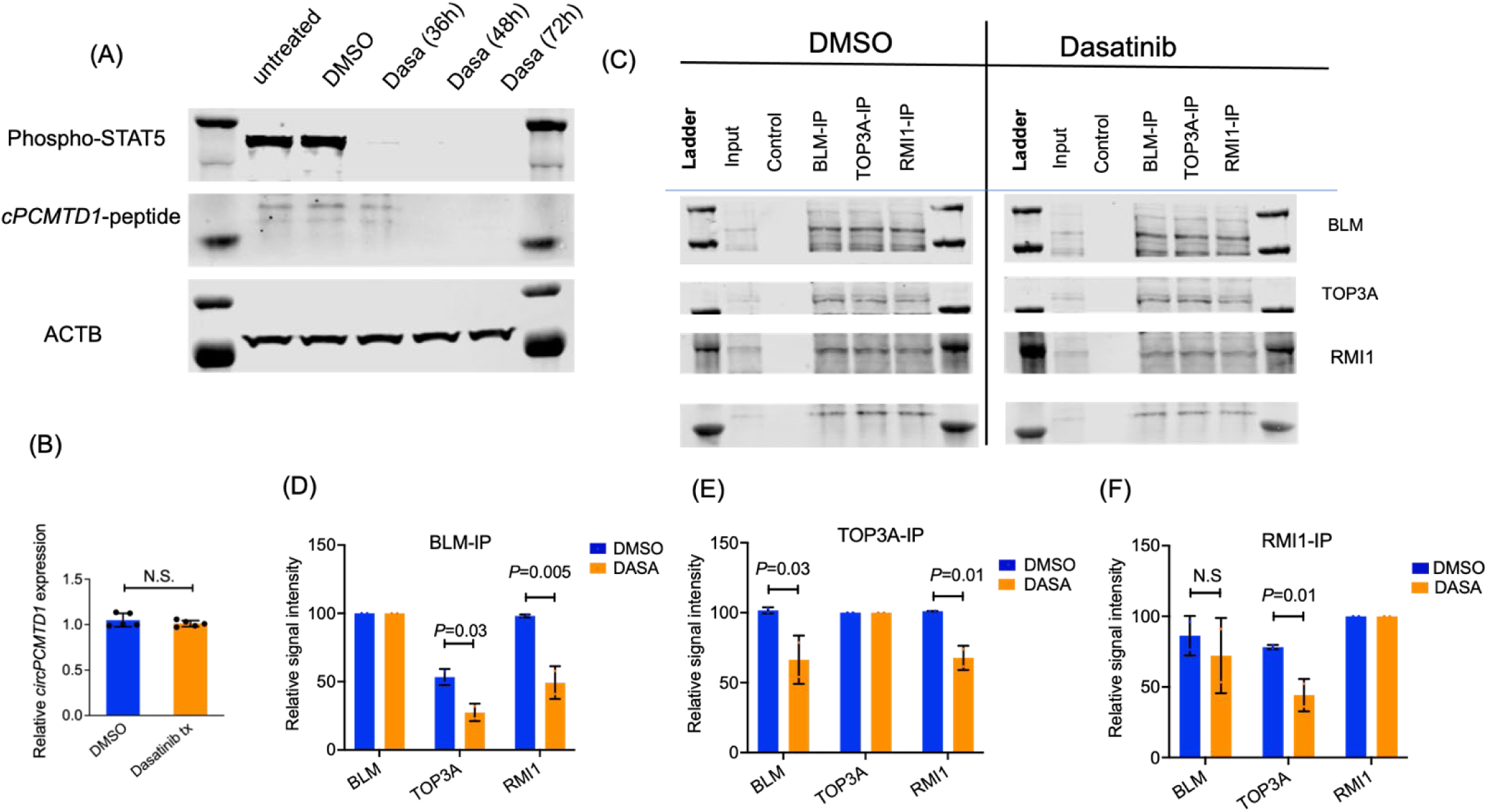
Blocking of the BCR/ABL signaling cascade reduces expression levels of the *circPCMTD1* derived peptide and impairs its interaction with the proteins of the BTR complex. **(A)** Western blot analysis showing phospho-STAT5 levels and *circPCMTD1*-derived peptide in K562 cells treated with Dasatinib at a 10 nM concentration at different timepoints (36, 48 and 72 hours). **(B)** Relative *circPCMTD1* RNA expression in K-562 cells following treatment with Dasatinib (1 nM concentration, 48h after treatment). (C) Coimmunoprecipitation experiments for the BTR complex following treatment with Dasatinib in K-562 cells. Cells were obtained 36h after initiation of treatment. Dasatinib was used at a 10 nM concentration. (D-F) Quantification of the strength of interaction among the BTR proteins following Dasatinib treatment (as depicted in C) by Image J. In (B), (D-F), P values were calculated using paired two-sided t-tests. In the figures, heights of bar graphs indicate mean values with standard deviation. Error bars indicate highest and lowest values in each population.

## DISCUSSION

CircRNAs are a novel class of RNA molecules which are gaining recognition for their functional role in health and disease.^1,2^ With regards to their role in myeloid malignancies, oncogenic circRNAs have been shown to stem from recurrent chromosomal translocations and be functionally relevant in AML with abnormal karyotypes.^22^ CircRNAs have also been shown to associate with outcome of CN-AML patients and to have biologic implications in this disease setting.^21,23^ A distinct circRNA-signature associated with *U2AF1* mutations in MDS has also been recently reported.^39^ In this work, we focus on the functional role of a previously uncharacterized circRNA, named *circPCMTD1*, in BCR/ABL-positive leukemias.

We first evaluated the effect of prognostic circRNA depletion in multiple leukemia cell lines. We detected a cell-type specific effect in the essentiality of *circPCMTD1*, as only leukemia cell lines positive for the *BCR/ABL1* translocation (i.e., K-562 and LAMA-84) were vulnerable to *circPCMTD1* depletion. We therefore focused on CML, which is typically associated with this genetic lesion. We performed targeted *circPCMTD1* profiling in CML patients in chronic phase, acceleration and blast crisis and found increased abundance of *circPCMTD1* in advanced disease stages, indicating a potential role of higher *circPCMTD1* expression in disease progression and aggressive phenotypes.

In experiments we conducted to characterize the mechanistic basis of *circPCMTD1* essentiality, we found that *circPCMTD1* depletion led to a notable activation of the DNA damage response pathway. Importantly, we observed increase of γH2AX-which is an established marker of DNA damage response-in leukemic cell lines and blasts of patients with *BCR/ABL1*-positive CML in blast crisis. In contrast (and in keeping with our observations in cell lines), *circPCMTD1* depletion did not affect H2Ax phosphorylation in blasts of AML patients.

Although predominantly associated with nuclear functions, we found that *circPCMTD1* was enriched in the cytoplasm and associated with the ribosomes of the leukemic blasts. Although RNA transcripts in circular configuration have been long reported to have a protein-coding potential,^34^ peptides derived from naturally occurring circRNAs have been recently identified as relevant for important cellular functions.^12–14^ The increased half-life of circRNAs, their resistance to exonuclease degradation and their potential for increased peptide output could underpin the biological advantages of encoding peptides and proteins in circularized transcripts.^35^ The common features of the peptide-producing circRNAs are the presence of a functional open reding frame in the circular sequence (i.e. with a start and an in-frame stop codon) as well as the capacity of the transcripts to interact with the translational machinery of the cells. In the case of *circPCMTD1*, the circular transcript hijacks the translation initiation codon in exon 2, which is in-frame with a stop codon in the 5-prime untranslated region of the linear *PCMTD1* transcript. We used a part of the peptide sequence which would be uniquely present in the predicted *circPCMTD1*-derived peptide and generated a custom-designed antibody. This antibody detected a protein band of approximately 30 KD, which decreased in intensity upon *circPCMTD1* knock-down.

To further characterize the mechanistic basis of *circPCMTD1* essentiality, we performed immunoprecipitation experiments followed by mass spectrometry. We found approximately 23 proteins that were candidate interactors of the *circPCMTD1*-derived peptide. Among the top scoring candidate interactors, we identified 3 proteins that are known to interact with each other and form the BTR complex (i.e., BLM, RMI1 and TOP3A). We validated the detected contacts with reciprocal immunoprecipitations and could further show that *circPCMTD1* depletion led to a decrease in the degree of interaction between the BML, RMI1 and TOP3A proteins and the formation of the BTR complex. We also confirmed that simultaneous depletion of the BTR complex proteins recapitulates the phenotype that is associated with *circPCMTD1* depletion.

The BTR complex plays a key role in genome integrity maintenance and DNA repair.^31^ Most notably, it is recruited to sites of DNA replication stress and facilitates the restarting of stalled replication forks.^31–33^ The chromosomal translocation of BCR/ABL has been associated with genomic instability and an increase in spontaneously occurring DNA damage.^36–38^ It can therefore be hypothesized that the increase in genotoxic stress that characterizes the CML-initiating event also increases the dependency of the leukemic cells on molecular mechanisms that facilitate DNA repair and increase tolerance to genomic stress. This dependency would be even more pronounced in the context of increased cell proliferation and cellular stress, during disease progression to accelerated phase and blast crisis. This link could provide the mechanistic basis for the sensitivity of BCR/ABL positive blasts to *circPCMTD1* depletion and subsequent aberrant activation of the DNA damage response.

To further dissect the mechanistic association between *circPCMTD1* and the presence of BCR/ABL1 kinase, we performed experiments with TKIs which block the constitutively active ABL1 kinase. We found that TKI treatment decreased the abundance of the *circPCMTD1*-derived peptide, without altering *circPCMTD1* expression levels. Notably, blocking of the BCR/ABL1 signaling cascade impaired the strength of interaction between the *circPCMTD1*-derived peptide and the BTR complex, indicating a direct link between the constitutively active tyrosine kinase and the ability of the *circPCMTD1*-derived peptide to associate with the BTR complex and regulate the DNA damage response.

The advent of TKIs has revolutionized the treatment of patients with CML. It is however true that a number of patients do not achieve complete eradication of their leukemic clones after treatment with frontline TKIs and are at risk of developing resistance to the targeted therapies.^40^ In addition, in the cases of patients who present with or later develop advanced stage disease (i.e., accelerated phase or blast crisis) TKI treatment is not sufficient to harness the leukemia and these patients require more aggressive therapeutic approaches.^40^ To evaluate the therapeutic utility of *circPCMTD1* targeting in advanced stage BCR/ABL-positive leukemias we performed experiments in preclinical animal models. We used blasts of CML patients in leukemic transformation to generate murine xenografts and could show that *in vivo* targeting of *circPCMTD1* led to a significant prolongation in the survival of the treated mice. Of note, approximately one third of the mice that were treated with *circPCMTD1*-targeting oligos had no residual leukemia when evaluated after completion of treatment.

In summary, herein we report on the functional significance of a previously uncharacterized circRNA, named *circPCMTD1* in acute leukemias. We identified a pronounced effect of *circPCMTD1* depletion in leukemic cells with *BCR/ABL1* translocations. Mechanistically we could show that *circPCMTD1* generated a peptide which interacted with the DNA damage response machinery and regulated the capacity of the leukemic blasts to tolerate genomic stress. Using preclinical models, we could show that *circPCMTD1*-targeting is a promising therapeutic strategy in the clinical setting of advanced stage *BCR/ABL1*-positive myeloid malignancies, for which TKI treatment is often inadequate and novel therapeutic approaches are needed.

## METHODS

### Patient samples

For the profiling of younger adult patients with CN-AML, pretreatment samples of patients enrolled on clinical trials of the Alliance for Clinical Trials in Oncology/Cancer and Leukemia Group B and treated with frontline chemotherapy regimens were analyzed as previously described.^21^

For RNA sequencing of CML patients, diagnosis samples of patients in chronic phase, accelerated phase or blast crises, as defined by established criteria,^40^ were analyzed with total RNA sequencing following ribosomal RNA depletion, using the Illumina 2500 device. For further details regarding library preparations and bioinformatic analyses please refer to the supplementary material.

### *In vitro* experiments

For *in vitro* experiments the following AML cell lines were purchased by biorepositories (i.e., ATCC, DKFZ): KG-1, THP-1, MOLM-13, K562, LAMA-84. The cell lines were cultured in RPMI media, supplemented with 10% of Fetal Bovine serum and 1% of antibiotics/antifungals and maintained at standard conditions (i.e., 37 degrees Celsius and 5% CO_2_, with saturating humidity). Patient samples were thawed and cultured in SFEM media, supplemented with BIT serum substitute (Stem Cell Technologies) and cytokines (SCF, FLT3-ligand, TPO, all purchased by Peprotech).

Delivery of locked nucleic acids targeting the back-splicing region of the *circPCMTD1* was performed via electroporation, using the Lonza electroporation platform and according to the instruction of the manufacturer. Solution L was used for the electroporation of K-562 and LAMA-84 cells and Monocyte solution was used for patient samples. Gapmers were transfected at a final concentration of 500 nM. The sequences of the gapmers that were used are provided in supplemental Table S5.

To confirm depletion of the targeted transcripts, RNA was extracted 24h after transfection using the Trizol reagent and standard protocols for chloroform-based RNA extraction. RNA was reverse transcribed to cDNA using random primers and the SuperScript III Kit according to the instruction of the manufacturer (Invitrogen).

### Cell cycle analysis

Cell cycle analysis was performed using the BrdU labeling kit, according to the instructions of the manufacturer (BD Pharmigen). In brief, 48h following delivery of the *circPCMTD1* targeting oligos, cells were labelled with BrdU for 1 hour, and were then fixed (incubation in Cytofix reagent) and permeabilized (Cytoperm reagent). Treated cells were stained with BrdU labelling antibodies and 7-AAD prior to analysis. Analysis was performed using standard flow cytometry experiments. Cell viability assays were performed using the annexin/PI staining protocol at 96h following delivery of gapmers.

### CyTOF experiments

Massively parallel flow cytometry experiments were conducted as previously described.^41^ In brief, cells treated with *circPCMTD1* gapmers versus controls were labelled with Ethynyldeoxyuridine (EdU) for 10 minutes, 48 hours after delivery of the targeting oligos. Cells were resuspended in CyFACS buffer, incubated with primary and secondary CyTOF antibody cocktails, washed and stained with intercalation buffers prior to being analyzed on the CyTOF device.

### RNA sequencing analysis

To evaluate the impact of *circPCMTD1* depletion on the transcriptome, RNA was extracted 48h post transfection with *circPCMTD1*-targeting gapmers. Libraries were generated using the ribo-zero depletion protocol (Illumina) by the core facility at Utah Medical center. Libraries were analyzed as outlined in the supplementary material.

### Intracellular staining for gH2Ax

The staining and flow acquisition were performed according to published protocols. Briefly, cells were permeabilized using the fixation/permeabilization solution and then stained with AlexaFluor 488-conjugated anti-γH2AX antibody, while DNA was counterstained with DAPI, according to the instruction of the manufacturer (BD Biosciences). Cells were analyzed by flow cytometry.

### DNA fiber assay

Cells were labeled with EdU (100 μM) for 30 min and then with BrdU (250 μM) for 30 min in the presence or absence of various inhibitors. Cells suspended in 1xPBS (2.5 μl of suspension at ∼10^6 cells/ml) were spotted onto a glass slide and allowed to dry. Cells were subsequently treated with 7.5 μl of spreading buffer (0.5% SDS, 200 mM Tris-HCl pH7.4, 50 mM EDTA) for 3-5 min. Slides were tilted (∼15°) to allow cell lysates to slowly run down the slide. The DNA on slides was air dried, fixed in methanol-acetic acid (3:1) for 2 min, and dried overnight at room temperature. Fixed DNA was denatured with 2.5 N HCl for 30 min at room temperature. Then, DNA was washed 3 times for 5 min with PBS-T (PBS + 0.05% Tween-20) and blocked in PBS-T containing 2% BSA for 30 min at 37°C. DNA fibers were incubated with rat anti-EdU (OBT0030, AbDSerotec) (1:200) and mouse anti-BrdU (BD Biosciences) (1:20) antibodies for 1 h at 37°C, followed by the incubation with Alexa-488-conjugated anti-mouse (1:100) and Cy3-conjugated anti-rat (Jackson ImmunoResearch) (1:100) secondary antibodies for 30 min 37°C. Slides were washed 3 times with PBS-T containing 1% BSA and mounted using VectaShield (Vector Labs). DNA fibers were imaged at 60X with a Nikon 90i microscope.

### Comet assays

For comet assays, K562 and LAMA-84 cells were treated with scramble versus *circPCMTD1* or *linPCMTD1*-targeting gapmers. Etoposide was used as DNA damaging agent (positive control). The R&D systems kit was used according to the instructions of the manufacturer. In brief, cells were embedded onto agarose and then loaded on slides. The slides were immersed into lysis solution, placed in alkaline buffer and run through electrophoresis. Cells are then stained with a fluorescent dye and visualized under a microscope. Comet assay slides were scored blindly (for presence of a tail) and data was averaged as average with SEM from three independent experiments. An average of 120 cells were analyzed for each experiment.

### Sucrose gradient based polysome profiling

For the isolation of polysome fractions and polysome profiling, 10×106 K562 cells were grown at a concentration of 5 × 105 cells/ml. Prior to lysis, a freshly prepared DMSO-solution of cycloheximide was added to the medium to a final concentration of 100 µg/ml. Cells were incubated for 10 min at 37 °C/5% CO_2_, washed in cycloheximide-containing PBS (100 µg/ml) and lysed in lysis buffer [(50 mM Tris-HCl pH 7.5, 10 mM KCl; 10 mM MgCl_2_, 150 mM NaCl, 2 mM DTT, 0.5 mM PMSF, 200 μg/l cycloheximide, 0.2% IGEPAL, and protease inhibitors: leupeptin (Sigma-Aldrich), pepstatin A (Sigma-Aldrich), and aprotinin (Sigma-Aldrich)]. Lysates were incubated on ice for 10 min and cleared by centrifugation in a tabletop microcentrifuge (16,000 × g, 10 min, 4 °C). The supernatants were collected into fresh tubes and flash frozen until polysome isolation or profiling was conducted.

Cytoplasmic extracts were thawed on ice. The equivalent of 4.5 million K562 cell lysates was layered onto each 11 ml of 10–50% linear sucrose gradient. Gradients were then centrifuged (35,000 rpm, 3 h, 4 °C) in a Sorvall-TH641 rotor at 4 °C. Gradient fractions of 500 uls each were collected consecutively with continuous measurement of UV absorbance at 254 nm.

For measurements of RNA transcripts, each isolated polysomal fraction was mixed with 1 ml of Trizol reagent and standard RNA extractions were performed. RNA samples were pooled together prior to reverse transcription and RT-qPCR. Cell pellets of 0.45 million cells were lysed with Trizol reagent and standard RNA extraction was performed. The unfractionated cells were used as input material, to determine the relative abundance of the transcripts on each isolated fraction of the sucrose gradient.

### In vivo studies

For in vivo studies, we used blasts of a CML patient in blast crisis and engrafted NOD.Cg-*Prkdc^scid^ Il2rgtm*^1W*jl*^/SzJ (NSG) mice. Prior to leukemic blast infusion, the animals were treated with busulfan [20 mg/kg weight with intraperitoneal (i.p.) injection]. On day 1 post-treatment the leukemic blasts were injected via the tail vein of the animals. Female mice of 6-8 weeks of age were used for the reported experiments. All animal studies were conducted according to protocols approved by the Institutional Animal Care and Use Committees of The Ohio State University (IRB protocol #2018C0072).

On day 10 post transplantation, NSG mice were assigned to groups to receive anti-*circPCMTD1* or non-targeting-control gapmers.

For the *in vivo* knock-down of *circPCMTD1*, *circPCMTD1*-targeting gapmers or non-targeting controls were packaged into cationic lipid nanoparticles (LNPs) conjugated with human transferrin, as previously described.^23^ In brief, the LNPs consisted of 1, 2-dioleoyl-3-trimethylammonium-propane (DOTAP) and 1,2-dioleoyl-sn-glycero-3-phosphocholine (DOPC) at a molar ratio of 1-to-1. DOTAP (18.8 mg) and DOPC (21.2 mg) were dissolved in 1 ml of ethanol and were then injected into 9 ml of HEPES buffer (20 mM, pH 7.4). After 5 min of incubation in a water bath sonicator, the nanoparticles were passed through a 0.22 µm sterile filter. Solutions of gapmers were mixed with lipids at a 1-to-10 weight ratio, vortexed and incubated at room temperature for 10 min. The lipid-oligo formulation was then mixed with a human transferrin solution. The mixtures were vortexed and sonicated in a water bath sonicator for 10 min and incubated for an hour at 37 °C. For the first week of treatment mice were treated with intravenous injections on days 1, 3, and 5 (of 5 mg of gapmers/kg weight) and i.p. injections on days 2 and 4 (of 10 mg of gapmers/kg weight). The mice received only i.p. injections (5/week) for the remainder of the treatment course (i.e., a total of three weeks of treatment).

### Western blotting, immunoprecipitation and mass spectrometry analysis

For Western blot analysis of the gH2Ax, CHK1, RPA-32, ATM, DNAPK and ATR proteins, manipulated cells were lysed directly with RIPA buffer. The cells were loaded onto gels and proteins were electrophoresed and separated based on molecular weight.

For the immunoprecipitation experiments, antibodies were cross-linked to A/G magnetic beads followed standard protocols provided by Abcam. In brief, the antibodies were incubated for an hour with A/G magnetic beads. The beads were washed x3 with immunoprecipitation buffer to remove excess antibody. The bead-antibody complexes were then incubated in for 30 mis in a Dimethyl palmitate solution at a final concentration of 6.5 mgs/ml (pH 8.5). Beads were washed with wash buffer x3 (0.2 M triethanolamine in PBS) and the cross-linking process was repeated twice. After this, the reaction was quenched with the addition of quenching buffer (50 mM ethanolamine in PBS) and beads were washed x3 in PBS. Beads with crosslinked antibodies were incubated with nuclear lysates for immunoprecipitation or mass spectrometry experiments.

To isolate nuclei 10e7 cells were harvested and resuspend in 2 mL of PBS, 2 ml of freshly prepared nuclear isolation buffer (1.28 M sucrose, 40 mM Tris-HCl pH 7.5, 20 mM MgCl_2_, 4% Triton X-100) and water (6 mL). Resuspended cells were kept on ice for 20 min (with frequent mixing). Nuclei were pelleted by centrifugation at 2,500 g for 15 min and resuspended in 1ml of RIP buffer (150 mM KCl, 25 mM Tris pH 7.4, 5 mM EDTA, 0.5 mM DTT, 0.5% NP40, Protease inhibitors). Nuclei were further sheared with a dounce homogenizer (15-20 strokes) on ice. Nuclear lysates were cleared off insoluble membranes and debris by centrifugation at 13,000 rpm for 10 min at 4 degrees. Nuclear lysates were incubated crosslinked antibody-bead conjugates at 4 degrees overnight. Beads were retained with magnets and captures were washed x3 in RIP buffer. Captured proteins were eluded by incubation in a glycine gradient (to a final concentration of 1 M).

### Mass Spectrometry Analysis

Proteins isolated following immunoprecipitation were incubated overnight with trichloroacetic acid (Ricca Chemical Company) and centrifugated (16,000 × *g*, 10 min, 4 °C). Precipitates were then digested with Trypsin (Promega) and Endopeptidase Lys-C (Wako) according to the instructions of the manufacturers, purified with HiPPR detergent removal columns (Life Technologies) and analyzed with liquid chromatography tandem mass spectrometry (LC/MSMS) on a Thermo Fusion Orbitrap Mass Spectrometer using 1 h LC gradient. Mass spectrometry analyses were performed in duplicates and the results were filtered for number and quality of identified peptides.

### Antibody Generation

Rabbit polyclonal antibodies were generated as a for-fee service by the Abclone company.

### Statistics and Reproducibility

Clinical and biological characteristics of patients were compared using the Fisher’s exact and Wilcoxon rank-sum tests for categorical and continuous variables, respectively. For CR, we calculated *P*-values using Fisher’s exact test. For time-to-event analyses, we calculated survival estimates using the Kaplan-Meier method and compared groups using the log-rank test. Analyses were preformed using SAS 9.4 and TIBCO Spotfire S+ 8.2.

For in vivo experiments, animal cohort sizes were not predetermined by statistical tests. Animals were divided into cohorts to ensure that weights and percentages of engrafted human blasts were similar among treatment groups. Researchers administering treatment or collecting data were not blinded to the conditions of the experiments. No data were excluded from the analyses. Statistical differences between survival rates were analyzed by comparison of Kaplan–Meier curves using the log-rank (Mantel–Cox) test.

Bar graphs and box plots are presented as mean ± s.e.m. For statistical comparisons between experimental groups, ANOVA (when multiple groups were compared simultaneously), followed by Mann–Whitney tests or paired t-tests were conducted. Statistical analyses were performed using Prism software (GraphPad version 10.0.0).

## ACKNOWLEDGEMENTS

R.B. was supported by a MERIT Award from the National Institutes of Health (NCI R37-CA252081) and a Research Scholar Grant from the American Cancer Society (RSG-24-1249960-01-DMC).

## AUTHOR CONTRIBUTIONS

D.P., A.P.U., and R.G. designed the study. D.P., A.P.U., R.B., R.K., and L.W performed laboratory experiments. D.P. performed proteomic analyses. L.W. performed polysome fractionation. A.P.U. performed animal studies. D.N and C.G. performed bioinformatics and biostatistical analyses. G.K.B. oversaw CyTOF experiments and analyzed data. A.P. and S.K. designed oligonucleotides/gapmers. D. P., K.M., A.-K.E., I.A., G.S. A.M.D. and R.G. oversaw experiments and data analysis. D.P., A.P.U. and R.G. prepared the manuscript with input from all co-authors.

## REFERENCES

1. Kristensen, L.S., et al. The biogenesis, biology and characterization of circular RNAs. Nat. Rev. Genet. 20, 675–691 (2019).

2. Chen, L.L. The expanding regulatory mechanisms and cellular functions of circular RNAs. Nat. Rev. Mol. Cell Biol. 21, 475–490 (2020).

3. Nigro, J.M., et al. Scrambled exons. Cell. 64, 607–613 (1991).

4. Cocquerelle, C., et al. Splicing with inverted order of exons occurs proximal to large introns. EMBO J. 11, 1095–1098 (1992).

5. Hansen, T.B., et al. Natural RNA circles function as efficient microRNA sponges. Nature. 495, 384–388 (2013).

6. Memczak, S., et al. Circular RNAs are a large class of animal RNAs with regulatory potency. Nature. 495, 333–338 (2013).

7. Szabo, L. & Salzman, J. Detecting circular RNAs: bioinformatic and experimental challenges. Nat. Rev. Genet. 17, 679–692 (2016).

8. Jeck, W.R. & Sharpless, N.E. Detecting and characterizing circular RNAs. Nat. Biotechnol. 32, 453–461 (2014).

9. Hansen, T.B. Improved circRNA identification by combining prediction algorithms. Front. Cell Dev. Biol. 6, 20 (2018).

10. Ashwal-Fluss, R., et al. circRNA biogenesis competes with pre-mRNA splicing. Mol. Cell 56, 55–66 (2014).

11. Li, Z., et al. Exon-intron circular RNAs regulate transcription in the nucleus. Nat. Struct. Mol. Biol. 22, 256–264 (2015).

12. Pamudurti, N.R., et al. Translation of circRNAs. Mol. Cell 66, 9–21 (2017).

13. Legnini, I., et al. Circ-ZNF609 is a circular RNA that can be translated and functions in myogenesis. Mol. Cell 66, 22–37 (2017).

14. Yang, Y., et al. Extensive translation of circular RNAs driven by N6-methyladenosine. Cell Res. 27, 626–641 (2017).

15. Kristensen, L.S., et al. The emerging roles of circRNAs in cancer and oncology. Nat. Rev. Clin. Oncol. 19, 188–206 (2022).

16. Qu, S., et al. The emerging landscape of circular RNA in life processes. RNA Biol. 14, 992–999 (2017).

18. Chen, S., et al. Widespread and functional RNA circularization in localized prostate cancer. Cell 176, 831–843 (2019).

19. Vo, J.N., et al. The landscape of circular RNA in cancer. Cell 176, 869–881 (2019).

20. Hanniford, D., et al. Epigenetic silencing of CDR1as drives IGF2BP3-mediated melanoma invasion and metastasis. Cancer Cell 37, 55–70 (2020).

21. Meng, S., et al. CircRNA: functions and properties of a novel potential biomarker for cancer. Mol. Cancer 16, 94 (2017).

22. Guarnerio, J., et al. Oncogenic role of fusion-circRNAs derived from cancer-associated chromosomal translocations. Cell 165, 289–302 (2016).

23. Papaioannou, D., et al. Clinical and functional significance of circular RNAs in cytogenetically normal AML. Blood Adv 4, 239–251 (2020).

24. Hirsch, S., et al. Circular RNAs of the nucleophosmin (NPM1) gene in acute myeloid leukemia. Haematologica 102, 2039–2047 (2017).

25. Gravel, S., Chapman, J.R., Magill, C. & Jackson, S.P. DNA helicases Sgs1 and BLM promote DNA double-strand break resection. Genes Dev. 22, 2767–2772 (2008).

26. Nimonkar, A. V. et al. BLM-DNA2-RPA-MRN and EXO1-BLM-RPA-MRN constitute two DNA end resection machineries for human DNA break repair. Genes Dev. 25, 350–362 (2011).

27. Karow, J.K., Chakraverty, R.K. & Hickson, I.D. The Bloom’s syndrome gene product is a 3′-5′ DNA helicase. J. Biol. Chem. 272, 30611–30614 (1997).

28. Döhner, H., et al. Diagnosis and management of AML in adults: 2022 recommendations from an international expert panel on behalf of the ELN. Blood 140, 1345–1377 (2022).

29. Pommier, Y., Nussenzweig, A., Takeda, S. & Austin, C. Human topoisomerases and their roles in genome stability and organization. Nat. Rev. Mol. Cell Biol. 23, 407–427 (2022).

30. Bizard, A.H. & Hickson, I.D. The many lives of type IA topoisomerases. J. Biol. Chem. 295, 7138– 7153 (2020).

31. Bizard, A.H. & Hickson, I.D. The dissolution of double Holliday junctions. Cold Spring Harb. Perspect. Biol. 6, a016477 (2014).

32. Lu, R., et al. The FANCM-BLM-TOP3A-RMI complex suppresses alternative lengthening of telomeres (ALT). Nat. Commun. 10, 2252 (2019).

33. Manthei, K.A. & Keck, J.L. The BLM dissolvasome in DNA replication and repair. Cell. Mol. Life Sci. 70, 4067–4068 (2013).

34. Chen, C.Y. & Sarnow P. Initiation of protein synthesis by the eukaryotic translational apparatus on circular RNAs. Science 268, 415–417 (1995).

35. Chen, R., et al. Engineering circular RNA for enhanced protein production. Nat Biotechnol. 41, 262–272 (2023).

36. Nowicki, M.O., et al. BCR/ABL oncogenic kinase promotes unfaithful repair of the reactive oxygen species-dependent DNA double-strand breaks. Blood 104, 3746–3753 (2004).

37. Slupianek, A., Nowicki, M.O., Koptyra, M. & Skorski, T. BCR/ABL modifies the kinetics and fidelity of DNA double-strand breaks repair in hematopoietic cells. DNA Repair (Amst). 5, 243–250 (2006).

38. Brady, N., Gaymes, T.J., Cheung, M., Mufti, G.J. & Rassool, F.V. Increased error-prone NHEJ activity in myeloid leukemias is associated with DNA damage at sites that recruit key nonhomologous end-joining proteins. Cancer Res. 63, 1798–1805 (2003).

39. Wedge, E., et al. Impact of U2AF1 mutations on circular RNA expression in myelodysplastic neoplasms. Leukemia 37, 1113–1125 (2023).

40. Hochhaus, A., et al. European LeukemiaNet 2020 recommendations for treating chronic myeloid leukemia. Leukemia 34, 966–984 (2020).

41. Behbehani, G.K., et al. Mass cytometric functional profiling of acute myeloid leukemia defines cell-cycle and immunophenotypic properties that correlate with known responses to therapy. Cancer Discov. 5, 988–1003 (2015).

