## Supplemental Information for "*circPCMTD1*: A protein-coding circular RNA that regulates DNA damage response in *BCR/ABL*-positive leukemias"

### Patient samples

AML samples that were analyzed for circRNA expression and association with outcome were collected as part of clinical trials of the Cancer and Leukemia Group B/Alliance for Clinical Trials. All patients were treated with frontline chemotherapy regimens, as previously described.[1]

For *in vitro* and *in vivo* functional experiments, we used leukemic samples of AML and CML patients obtained after leukapheresis or diagnostic bone marrow biopsies and deposited in the Leukemia Tissue Bank of The Ohio State University. All patient samples were de-identified and stored under codes in a process compliant with the Health Insurance Portability and Accountability Act. All patients had provided written informed consent for the use of their biologic specimens for research purposes in alignment the Declaration of Helsinki. All experiments conducted with human patient samples were in compliance with the Institutional Review Board (IRB) of each participating institution.

### Transcriptome analysis: library generation, sequencing and data analysis

Extracted total RNA was assessed for quality on an Agilent 2100 Bioanalyzer (BioA) using the RNA 6000 Nanochip and for quantity on a Qubit 2.0 Fluorometer (Agilent Technologies, Santa Clara, CA) using the RNA HS Assay Kit. Samples with an RNA Integrity Number (RIN) greater than four, with no visible sign of genomic DNA (gDNA) contamination and a concentration of >40 ng/μL were used for total RNA library generation. RNA-seq libraries were prepared using the Illumina TruSeq Stranded Total

RNA Sample Prep Kit with RiboZero Gold (#RS1222201) according to the manufacturer's instructions. Sequencing was performed with the Illumina HiSeq 2500 system using the HiSeq version 3 sequencing reagents to an approximate cluster density of 800,000/mm$^2$. Image analysis, base calling, error estimation, and quality thresholds were performed using the HiSeq Controller Software (version 2.2.38) and the Real Time Analyzer software (version 1.18.64).

Transcript abundance was quantified using kallisto[2] (https://pachterlab.github.io/kallisto). The Genome Reference Consortium version 38 of the human genome was used as reference for the analysis.

**Definition of clinical endpoints and disease classification**

For patients with AML, clinical endpoints were defined according to generally accepted criteria.[3] Complete remission (CR) required a BM aspirate with cellularity >20% with maturation of all cell lines, <5% blasts and undetectable Auer rods, as well as an absolute neutrophil count of ≥1.5 x $10^9$/L, platelet count of >100 x $10^9$/L, and absence of leukemic blasts in peripheral blood. Evidence of extramedullary leukemia were evaluated and needed be ruled out to diagnose CR. All of the above had to persist for ≥4 weeks for CR to be determined.[3] Relapse was defined by the presence of ≥5% BM blasts, or circulating leukemic blasts, or the development of extramedullary leukemia. Disease-free survival (DFS) was measured from the date of CR until the date of relapse or death (from any cause); patients alive and in continuous first CR were censored at last follow-up. Overall survival (OS) was measured from the date of study entry until the date of death (from any cause); patients alive at last follow-up were censored.

Regarding the dataset of patients with chronic myeloid leukemia, de-identified samples were obtained and analyzed with real-time quantitative PCR. Presence of BCR-ABL translocation was confirmed with cytogenetic analysis and/or fluorescent in situ hybridization. As per published guidelines,[4,5] patients were diagnosed with CML in chronic phase if BCR/ABL was detected and myeloblasts were below 10% in blood or bone marrow. Patients with 10-19% blasts in blood or bone marrow, ≥20% basophils in blood, persistent thrombocytopenia (<100 × $10^9$/L) unrelated to therapy, thrombocytosis (platelet count >1000 × $10^9$/L) unresponsive to therapy, increasing spleen size and increasing WBC count unresponsive to therapy, or cytogenetic evidence of clonal evolution were classified as CML in accelerated phase. Blast percentages higher than 20% in blood or bone marrow, extramedullary blast proliferation, or large foci or clusters of blasts in the bone marrow biopsy defined the blast crisis stage.[4,5]

**Transcriptome analysis of leukemic blasts following *circPCMTD1* depletion**

To analyze RNA sequencing data following depletion of *circPCMTD1*, libraries were generated as described above. 48 hours post knockdown of *circPCMTD*1 in K562 cells, RNA was extracted from cells and submitted for bulk RNA-sequencing. Paired Fastq files were trimmed to remove adapter sequences using Trimgalore, and trimmed fastq files were aligned to the reference human genome GRCh38 using STAR tool. Gene counts were generated from aligned files using featureCounts. Differential gene expression analysis was done using DESeq2. Volcano plot shows significantly differentially expressed genes (adjusted P<0.05, Fold change cutoff +/- 1.5). GSEA analysis was performed to identify enriched biological processes.

### SUPPLEMENTAL REFERENCES


1. Papaioannou D, Volinia S, Nicolet D, et al. Clinical and functional significance of circular RNAs in cy-togenetically normal AML. *Blood Adv* 2020;4(2):239-251.
2. Bray NL, Pimentel H, Melsted P, Pachter L. Near-optimal probabilistic RNA-seq quantification. *Nat Biotechnol*. 2016;34:525-527.
3. Cheson BD, Cassileth PA, Head DR, et al. Report of the National Cancer Institute-sponsored workshop on definitions of diagnosis and response in acute myeloid leukemia. *J Clin Oncol*. 1990;8:813-819.
4. Lahaye T, Riehm B, Berger U, et al. Response and resistance in 300 patients with BCR-ABL-positive leukemias treated with imatinib in a single center: a 4.5-year follow-up. *Cancer* 2005;103:1659-1669.
5. Baccarani M, Saglio G, Goldman J, et al. Evolving concepts in the management of chronic myeloid leukemia: recommendations from an expert panel on behalf of the European LeukemiaNet. Blood. 2006;108:1809–20

### SUPPLEMENTAL TABLES

**Supplemental Table S1. Clinical Outcome of Younger Adult Patients With Cytogenetically Normal Acute Myeloid Leukemia by *circPCMTD1* expression status.**

| n=365 | Low c*PCMTD1** (n=182) | High c*PCMTD1** (n=183) | P† |
|---|---|---|---|
| CR, no. (%) | 98 (77) | 109 (86) | .11 |
| Disease-Free Survival (DFS) | | | .001 |
| Median (years) | 1.3 | 3.2 | |
| %Disease-free at 3 years | 29 (20-38) | 50 (40-59) | |
| %Disease-free at 5 years | 26 (18-35) | 47 (38-56) | |
| Overall Survival (OS) | | | .003 |
| Median (years) | 1.7 | 5.5 | |
| %Alive at 3 years | 40 (32-49) | 58 (49-66) | |
| %Alive at 5 years | 34 (26-42) | 50 (41-59) | |

* Median expression value was used as the cut point

†P-values for categorical variables are from Fisher's exact test, p-values for the time to event variables are from the log-rank test.

The median follow-up for those alive is 8.4 years, range: 0.6-21.2 years (n=97).
The median follow-up for those who have not had an event is 8.3 years, range: 0.6-21.2 years (n=70).

**Supplemental Table S2. Clinical and Molecular Features of Younger Adult Patients With Cytogenetically Normal Acute Myeloid Leukemia by *circPCMTD1* expression status.**

| **n=365** | **Low c*PCMTD1** (n=182)*** | **High c*PCMTD1** (n=183)*** | **P[§]** |
|---|---|---|---|
| Age (years) | | | .34 |
| Median | 46 | 46 | |
| Sex, no.(%) | | | .35 |
| Male | 99 (54) | 89 (49) | |
| Female | 84 (46) | 93 (51) | |
| Race, no.(%) | | | .10 |
| White | 157 (88) | 167 (93) | |
| Non-white | 22 (12) | 12 (7) | |
| Hemoglobin (g/dL) | | | **.003** |
| Median | 8.9 | 9.4 | |
| Platelet count (x109/L) | | | .37 |
| Median | 56 | 57 | |
| WBC count (x109/L) | | | **< .001** |
| Median | 39.9 | 23.7 | |
| %Blood Blasts | | | .33 |
| Median | 61 | 58 | |
| %Bone Marrow Blasts | | | **.004** |
| Median | 72 | 64 | |
| Extramedullary Involvement, no. (%) | | | .13 |
| Present | 60 (34) | 46 (26) | |
| Absent | 118 (66) | 132 (74) | |
| ELN Genetic Group, no. (%) | | | .19 |
| Favorable | 89 (50) | 103 (60) | |
| Intermediate | 53 (30) | 39 (23) | |
| Adverse | 35 (20) | 31 (18) | |
| *ASXL1*, no. (%) | | | **.04** |
| Mutated | 2 (1) | 10 (6) | |
| Wild-type | 169 (99) | 165 (94) | |
| *CEBPA*, no. (%) | | | **.03** |
| bZip Mutated | 19 (11) | 35 (20) | |
| Wild-type | 154 (89) | 140 (80) | |

* Median was used as the cut point

§P-values for categorical variables are from Fisher's exact test, P-values for continuous

**Supplemental Table S3. Differential Gene Expression following delivery of *circPCMTD1*-targeting oligos in BCR/ABL-positive leukemic cells (provided as Excel sheet).**

**Supplemental Table S4. Predicted nucleotide and amino acid sequence of the open reading frame contained in the *circPCMTD1* transcript (provided as Excel sheet).**

**Supplemental Table S5. Candidate protein interactors of the *circPCMTD1*-derived peptide as identified by immunoprecipitation experiments followed by Mass spectrometry (provided as Excel sheet).**

**Supplemental Table S6. Sequences of LNA-modified oligonucleotides designed for knockdown of circular RNA transcripts (provided as Excel sheet).**

**Supplemental Figure S1. Clinical Outcome of Younger Adult Patients With Cytogenetically Normal Acute Myeloid Leukemia by *circPCMTD1* expression status.** (A) Disease-free survival (DFS), (B) overall survival (OS).

**A** **B**

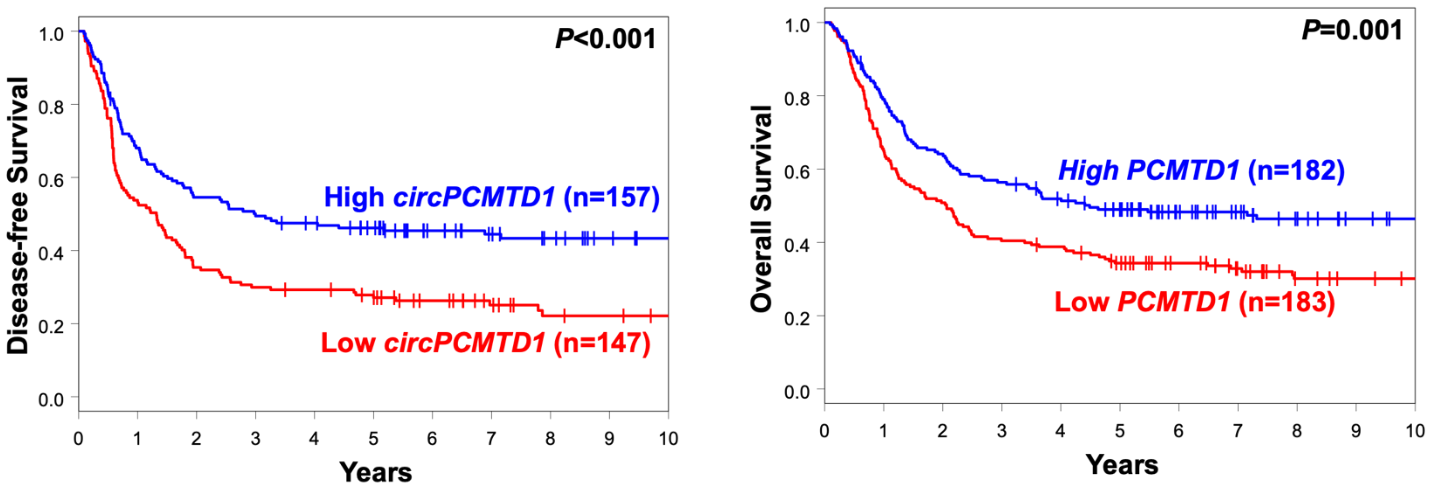

**Supplemental Figure S2. *CircPCMTD1* targeting does not affect cell cycle progression in acute leukemia cell lines without the BCR-ABL translocation** (A-D) BrDU labelling followed by cell cycle analysis in MOLM13 (A-C) and THP-1 (D-F) cells treated with scramble (A,D) versus *circPCMTD1* KD (B,E). (C) and (F) depict the results of three independent experiments in aggregate. Significance was tested using paired two-sided t-tests

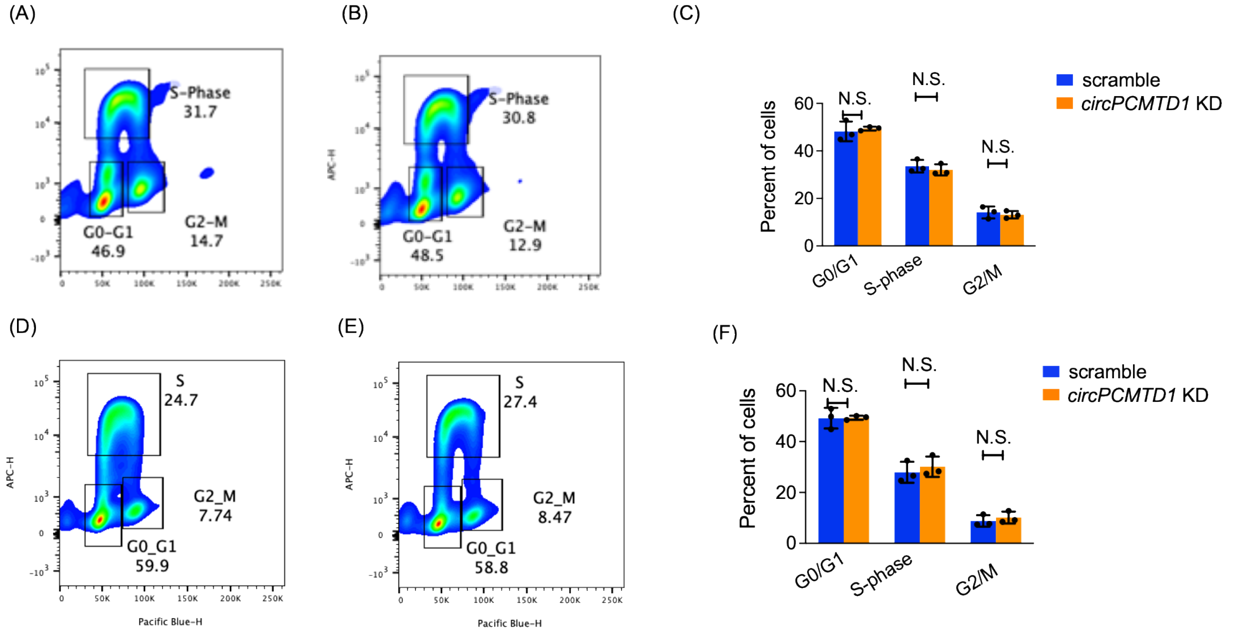

**Supplemental Figure S3. Targeting of the linear *PCMTD*1 transcript does not affect cell cycle progression in acute leukemia cell lines which harbor the BCR-ABL translocation** (A-B) BrDU labelling followed by cell cycle analysis in K-562 cells treated with scramble versus *linPCMTD1* KD. (C) depicts the results of three independent experiments in aggregate. Significance was tested with paired two-sided t-tests.

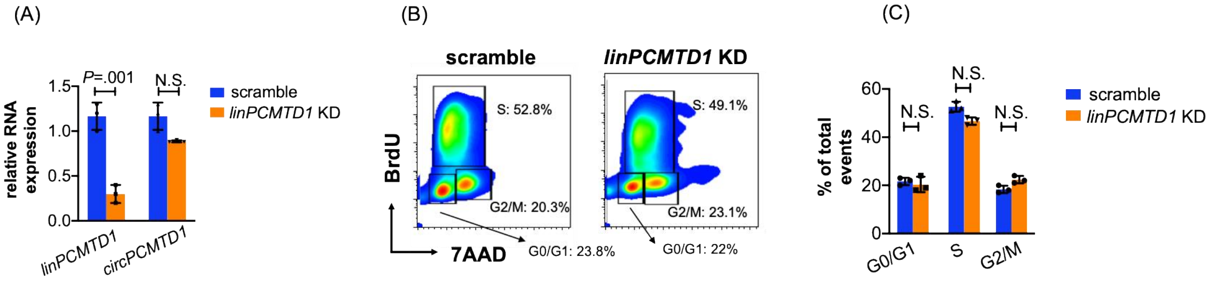

**Supplemental Figure S4. *CircPCMTD1* depletion activates the aberrant DNA damaging response in *BCR-ABL* positive leukemic blasts.** (A, B) Intracellular staining for phosphor-H2Ax levels followed by flow cytometry analysis in K-562 cells treated with (A) scramble versus (B) *circPCMTD1* KD. (C) depicts the results of three independent experiments in aggregate. Significance was tested with paired two-sided t-tests.

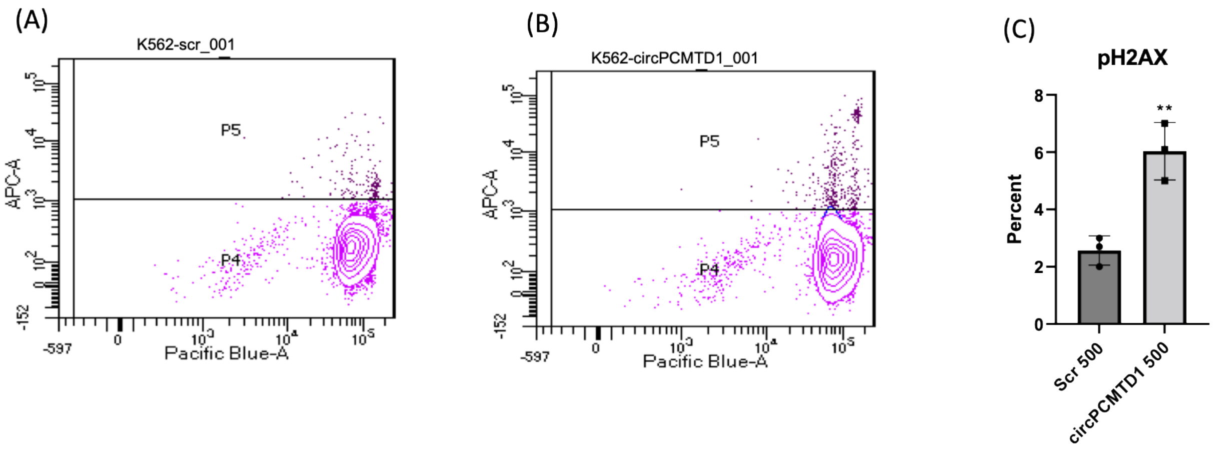

**Supplemental Figure S5. *CircPCMTD1* depletion does not affect levels of the BTR complex proteins.** Western blotting for the BLM, TOP3A, RMI1, ACTB and the circPCMTD1-derived peptide in (A) K-562 and (B) LAMA-84 cells, treated with scramble versus *circPCMTD1* KD at 24 and 48 hours post-delivery of *circPCMTD1*-targeting gapmers. Experiments were performed in duplicates. Results of one representative experiment are shown.

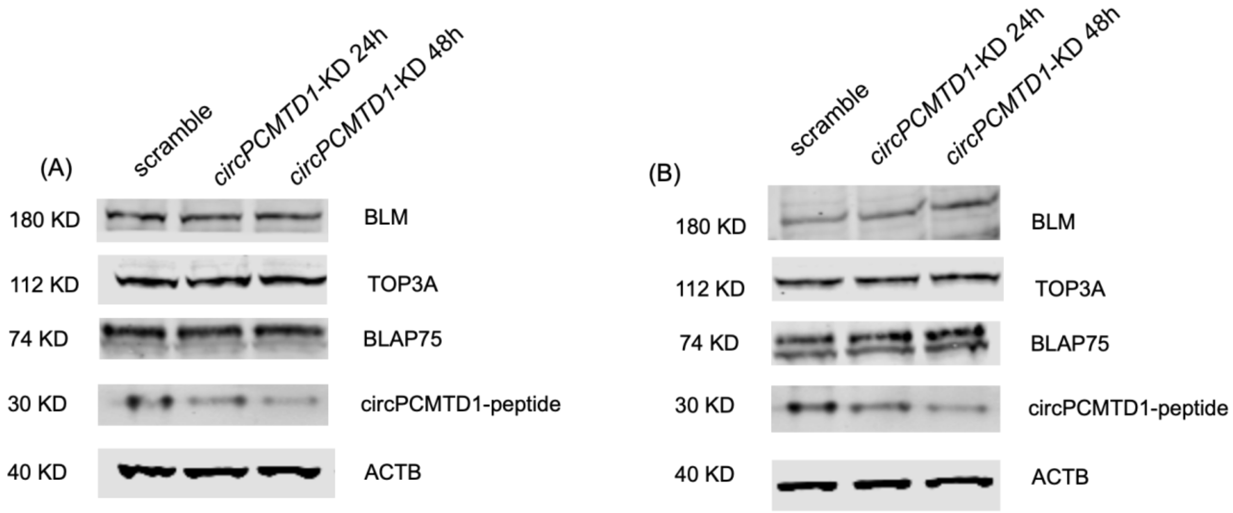

**Supplemental Figure S6: The *circPCMTD1*-derived peptide regulates the strength of the interaction among the proteins of the BTR complex in LAMA-84 cells.** (A) Immunoprecipitations with antibodies targeting the BLM, TOP3A, RMI1 proteins, followed by western blotting in LAMA-84 cells treated with scramble versus *circPCMTD1* KD. (B-D) Quantification of the strength of interaction among the BTR proteins following *circPCMTD1*-KD by Image J

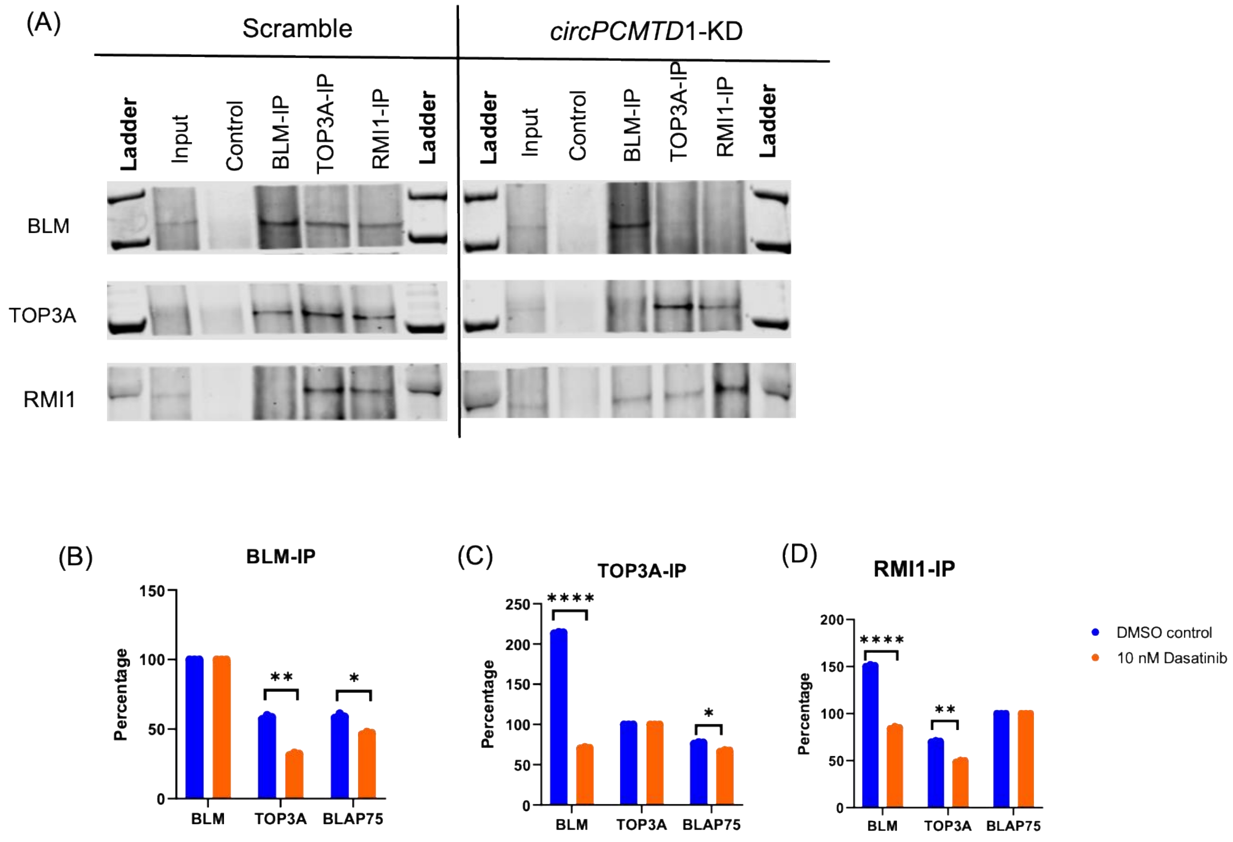

**Supplemental Figure S7: Knock-down of the BLM, TOP3A and RMI1 predominantly impacts on cell viability of BCR/ABL-positive leukemic blasts.** (A-C) RNA abundance of *BLM*, *TOP3A* and *RMI1* following delivery of respective antisense oligonucleotides in K562 cells. (D-F) BrdU-based cell cycle analysis of K-562 cells following delivery of *BLM*, *TOP3A* and *RMI1*-targeting oligonucleotides. Results of three independent experiments are depicted in aggregate. (H-I) Annexin-PI-based viability analysis of K-562 cells following delivery of *BLM*, *TOP3A* and *RMI1*-targeting oligonucleotides. Results of three independent experiments are depicted in aggregate.

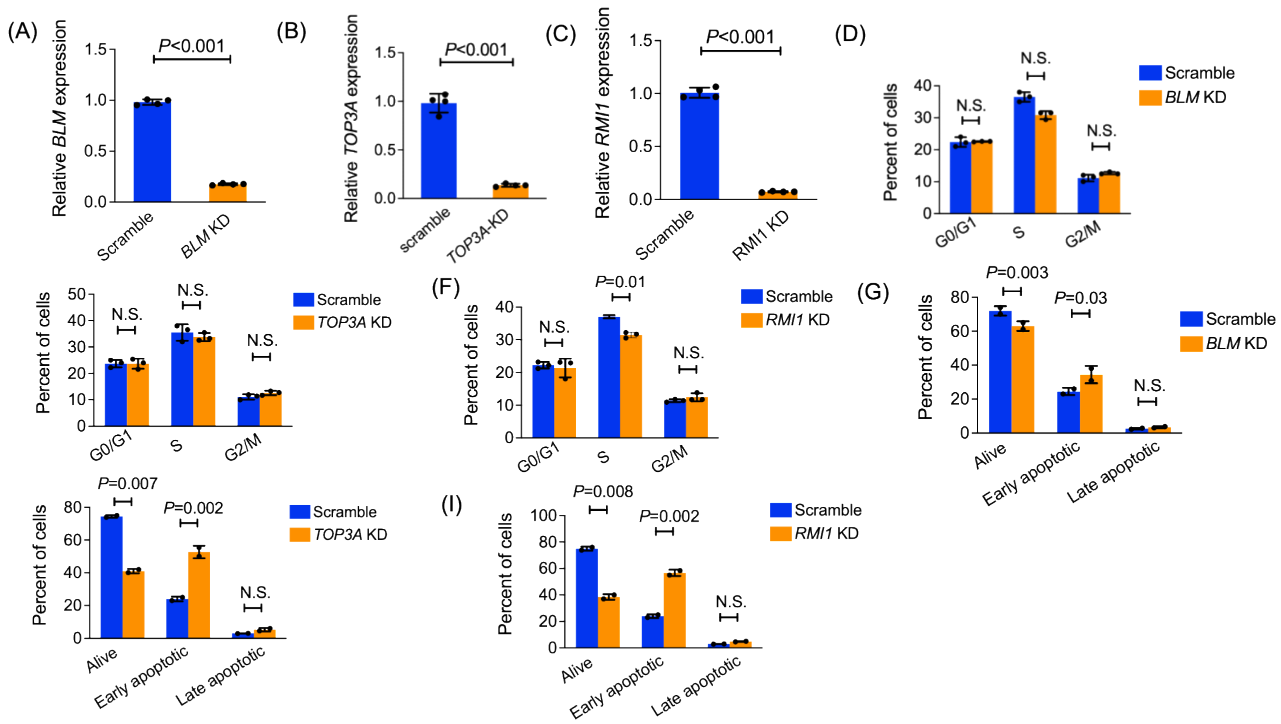
